# Efficacy and Safety of Immunosuppressive Treatment in IgA Nephropathy: A Meta-analysis of Randomized Controlled Trials

**DOI:** 10.1101/515023

**Authors:** Zheng Zhang, Yue Yang, Shi-min Jiang, Wen-ge Li

**Author notes:** **Corresponding author’s contact information:** Wen-ge Li, E- Mail.

## Abstract

**Background:** There is some controversy regarding the efficacy and safety of immunosuppressive agents for the treatment of kidney diseases. The recent STOP-IgAN and TESTING studies have focused attention on the application of immunosuppressive agents in IgA nephropathy (IgAN). This study investigated the benefits and risks of immunosuppressive agents in IgAN.

**Methods:** MEDLINE, EMBASE, the Cochrane Library, and article reference lists were searched for randomized controlled trials (RCTs) comparing immunosuppressive agents with any other non-immunosuppressive agents for treating IgAN. A meta-analysis was performed on the outcomes of proteinuria, creatinine (Cr), estimated glomerular filtration rate (eGFR), and adverse events in patients with IgAN, and trial sequential analyses were also performed for outcomes.

**Results:** Twenty-nine RCTs (1957 patients) that met our inclusion criteria were identified. Steroids (weighted mean difference [WMD] −0.70, 95% confidence interval [CI] −1.2 to −0.20), non-steroidal immunosuppressive agents (NSI) (WMD −0. 43, 95% CI −0.55 to −0.31), and combined steroidal and non-steroidal immunosuppressive agents (S&NSI) (WMD −1.46, 95% CI −2.13 to −0.79) therapy significantly reduced proteinuria levels in patients with IgAN. Steroid treatment significantly reduced the risk for end-stage renal disease (ESRD) (relative risk [RR] 0.39, CI 0.19 to 0.79). The immunosuppressive therapy group showed significant increases in gastrointestinal, hematological, dermatological, and genitourinary side effects, as well as impaired glucose tolerance or diabetes. Hyperkalemia was more common in the control group.

**Conclusion:** Immunosuppressive therapy can significantly reduce proteinuria and ESRD risk in patients with IgAN, but with a concomitant increase in adverse reactions. Therefore, care is required in the application of immunosuppressive agents in IgAN.

## Introduction

IgA nephropathy (IgAN) is one of the most common primary glomerular diseases (1). A systematic review demonstrated an overall population incidence of IgAN of 2.5/100000/year (2). There is still no uniform standard of treatment for IgAN. The 2012 Kidney Disease: Improving Global Outcomes (KDIGO) guidelines (3) for IgAN recommend treatment with a renin-angiotensin system (RAS) blocker, such as angiotensin-converting enzyme inhibitors (ACEIs) and angiotensin II receptor blockers (ARBs), in patients with proteinuria with protein excretion > 1 g/day. Corticosteroid therapy can be considered in patients with proteinuria > 1 g/day after 3–6 months of best supportive treatment and without renal failure. Intensive immunosuppression is reserved for patients with crescents in more than half the glomeruli and a rapid decline in renal function.

The publication of the Supportive versus Immunosuppressive Therapy of Progressive IgA Nephropathy (STOP-IgAN) trial in 2015 and Therapeutic Evaluation of Steroids in IgA Nephropathy Global (TESTING) trial in 2017 focused attention on the treatment of IgAN with immunosuppressive agents. According to the results of these two large randomized controlled trials (RCTs), there is still no clear evidence that immunosuppressive therapy can improve the prognosis of IgAN. Therefore, we retrieved RCTs on immunosuppressive therapy for IgAN, and performed a meta-analysis of the efficacy and safety of immunosuppressive therapy in this disease.

Immunosuppressive agents were divided into three subgroups for this meta-analysis: steroids, non-steroidal immunosuppressive (NSI) agents, and steroids combined with non-steroidal immunosuppressive (S&NSI) agents. Their efficacy and safety were compared relative to controls for the treatment of IgAN.

This meta-analysis was performed in accordance with the recommendations of the Cochrane handbook for systematic reviews of interventions (4) and is reported in compliance with the Preferred Reporting Items for Systematic Reviews and Meta-Analyses (PRISMA) statement guidelines (5). The protocol and registration information are available at http://www.crd.york.ac.uk/PROSPERO/ (CRD42018096197).

### Inclusion and exclusion criteria

This investigation required studies to meet the following inclusion criteria: the study was an RCT; the study compared different immunosuppressive agents versus non-immunosuppressive agents/placebo/no treatment; and study subjects were adult or pediatric patients with biopsy-proven IgAN.

Studies were rejected according to the following exclusion criteria: immunosuppressant not given orally or intravenously; study subjects with secondary IgAN; no data available for this study in the article, data included in other articles, or data repeated in other articles; and article not in English.

### Data sources and searches

The MEDLINE, EMBASE, and Cochrane Library medical databases were searched to retrieve relevant studies. Searches were performed in English, and each search retrieved studies that were published between establishment of the database and May 2018.

A comprehensive search strategy was established to ensure the comprehensive and accurate retrieval of studies. Specifically, the MEDLINE and Cochrane Library databases were searched using the method described in the Cochrane Policy Manual for optimizing the sensitivity and precision of the search process (6), whereas EMBASE was searched using a sensitivity–specificity filter optimized by the McMaster/Hedges team (7). The following search terms were used: IgAN, steroids, glucocorticoids, immunosuppressive agents, angiotensin-converting enzyme inhibitors, angiotensin receptor antagonists, and placebo. After completing the electronic query of the aforementioned databases, we also searched relevant professional journals manually.

### Data extraction and quality assessment

Two investigators (ZZ and YY) independently selected studies from the retrieved literature based on the inclusion criteria and extracted the data and analytical results of these studies. If the two investigators had different opinions regarding the quality of a study, a third investigator (SMJ) examined the disputed study and discussed it with the two aforementioned reviewers. Data were included for consideration only if discussions allowed the three authors to achieve consensus regarding the data.

If necessary, daily proteinuria was recalculated as g/day. Values for eGFR were based on the data provided by the authors of the included studies.

We evaluated treatment-related changes based on changes between the pre-treatment and post-treatment mean values and standard deviations (SDs) of the examined outcome measures. As the standard error of the mean (SEM) was used in some studies, we calculated the SD using the formula: SEM × square root of sample size. In addition, 95% confidence intervals (CIs) were used in some studies; we calculated the SD using the formula: ((upper limit of 95% CI – lower limit of 95% CI)/(2 × 1.96)) × √(n). Publication bias is defined as a condition in which studies with positive results are more likely to be published. Assessment of the risk of bias was performed following the Cochrane handbook.

### Risk of bias assessment

Two authors (ZZ and YY) independently assessed risk of bias using the Cochrane risk-of-bias tool (8). They reviewed each trial and gave a score of high, low, or unclear risk of bias according to the following criteria: random sequence generation, allocation concealment, blinding of participants and personnel to the study protocol, blinding of outcome assessment, incomplete outcome data, selective reporting, and other bias.

### Statistical analyses

To compare the effects of immunosuppressive agents and control treatment on proteinuria excretion and serum levels of creatinine, data on eGFR and end-stage renal disease (ESRD) were extracted for meta-analyses. Subgroup analyses were performed for each outcome based on the type of immunosuppressive agent.

For continuous outcomes, the differences in means and the 95% CI in mean change between baseline and end of treatment value were calculated for individual trials, and the weighted mean difference (WMD) was used as a summary estimator. Dichotomous outcome data from individual trials were analyzed using the relative risk (RR) measure and 95% CI. Heterogeneity of treatment effects between studies was investigated visually by examination of plots and statistically using the heterogeneity χ^2^ and I^2^ statistics. In all analyses, *P* < 0.05 was taken to indicate statistical significance. The fixed-effects and random-effects models were used for the meta-analysis of each indicator. Analyses were performed using Review Manager 5.2 (RevMan; Cochrane Collaboration, Oxford, UK).

### Trial sequential analyses

To evaluate whether the present meta-analysis had a sufficient sample size to reach firm conclusions about the effects of interventions, we performed trial sequential analyses (TSAs) for outcomes, which involves a cumulative meta-analysis to create a Z curve of the summarized observed effect (the cumulative number of included patients and events) and the monitoring boundaries for benefit and harm and estimate the optimal sample size (9). When the cumulative z curve crosses the trial sequential monitoring boundary, a sufficient level of evidence for the anticipated intervention effect may have been reached. If the z curve crosses none of the boundaries and the required information size has not been reached, there is insufficient evidence to reach a conclusion. These analyses were performed using the software TSA version 0.9 Beta (Copenhagen Trial Unit, Copenhagen, Denmark).

### Basic information regarding the included studies

After performing electronic and manual searches, 4,016 potentially relevant papers were obtained. After removing duplicated papers, 2,639 papers remained. After browsing the titles and abstracts, 53 papers were selected. After reading the entire text of these 53 papers, 24 papers were excluded, and 29 papers describing 25 trials with a total of 1957 patients were ultimately included. The literature selection process is illustrated in Figure 1, and detailed information regarding the examined studies is provided in Table 1 (10–38).

**Fig. 1.**
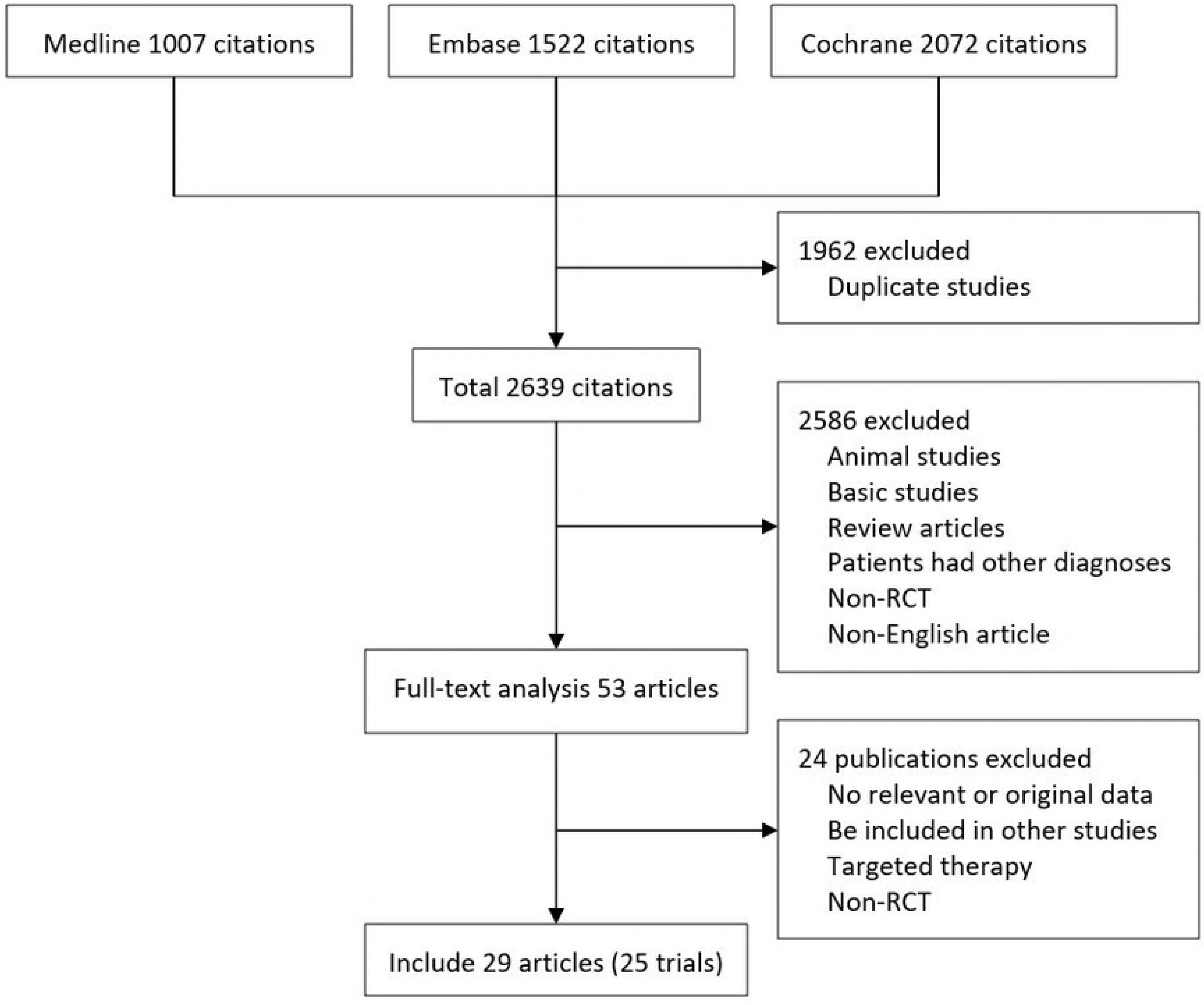
Results of systematic literature search on immunosuppressive treatment for IgAN.

**Table 1.**
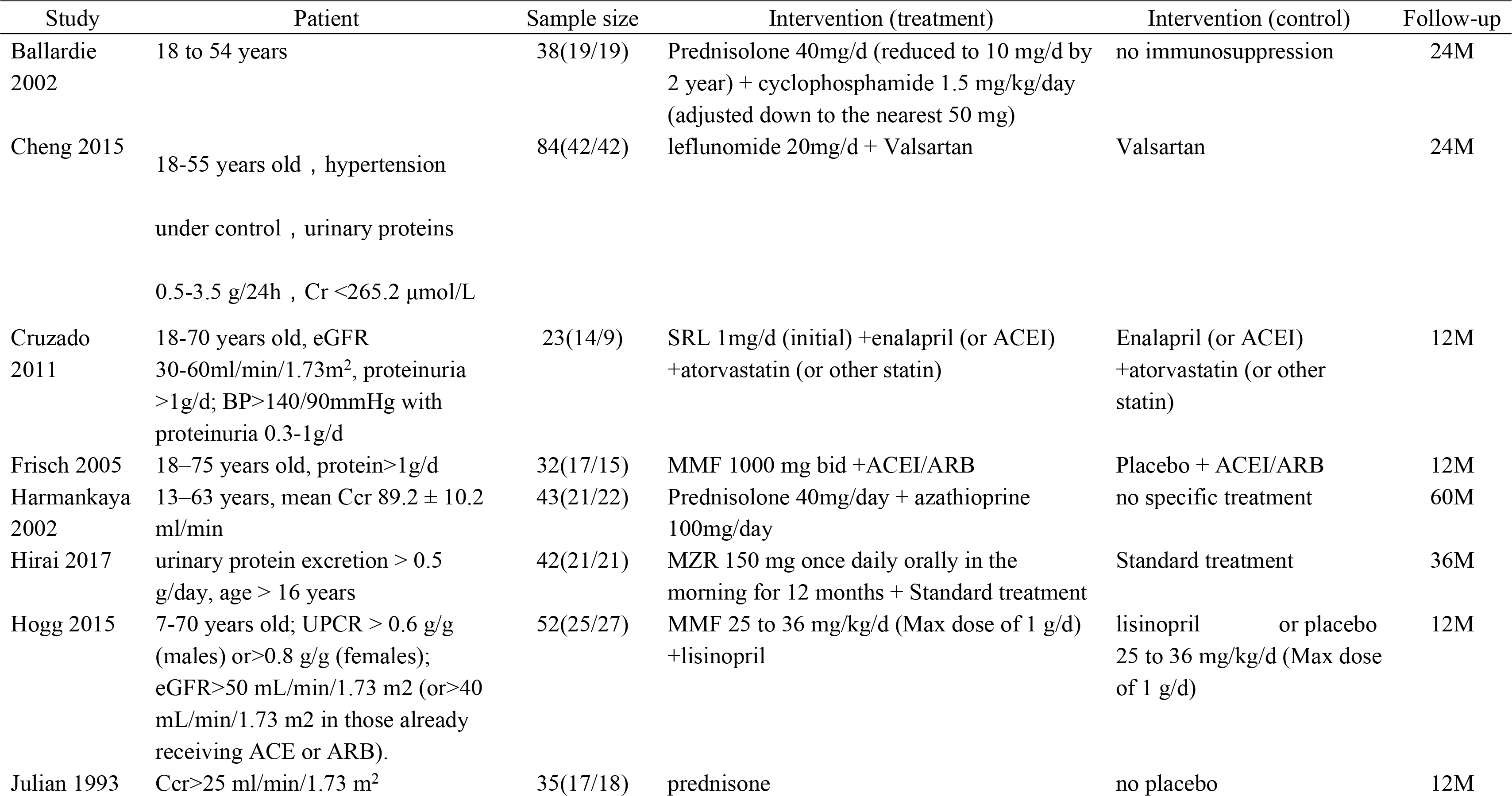
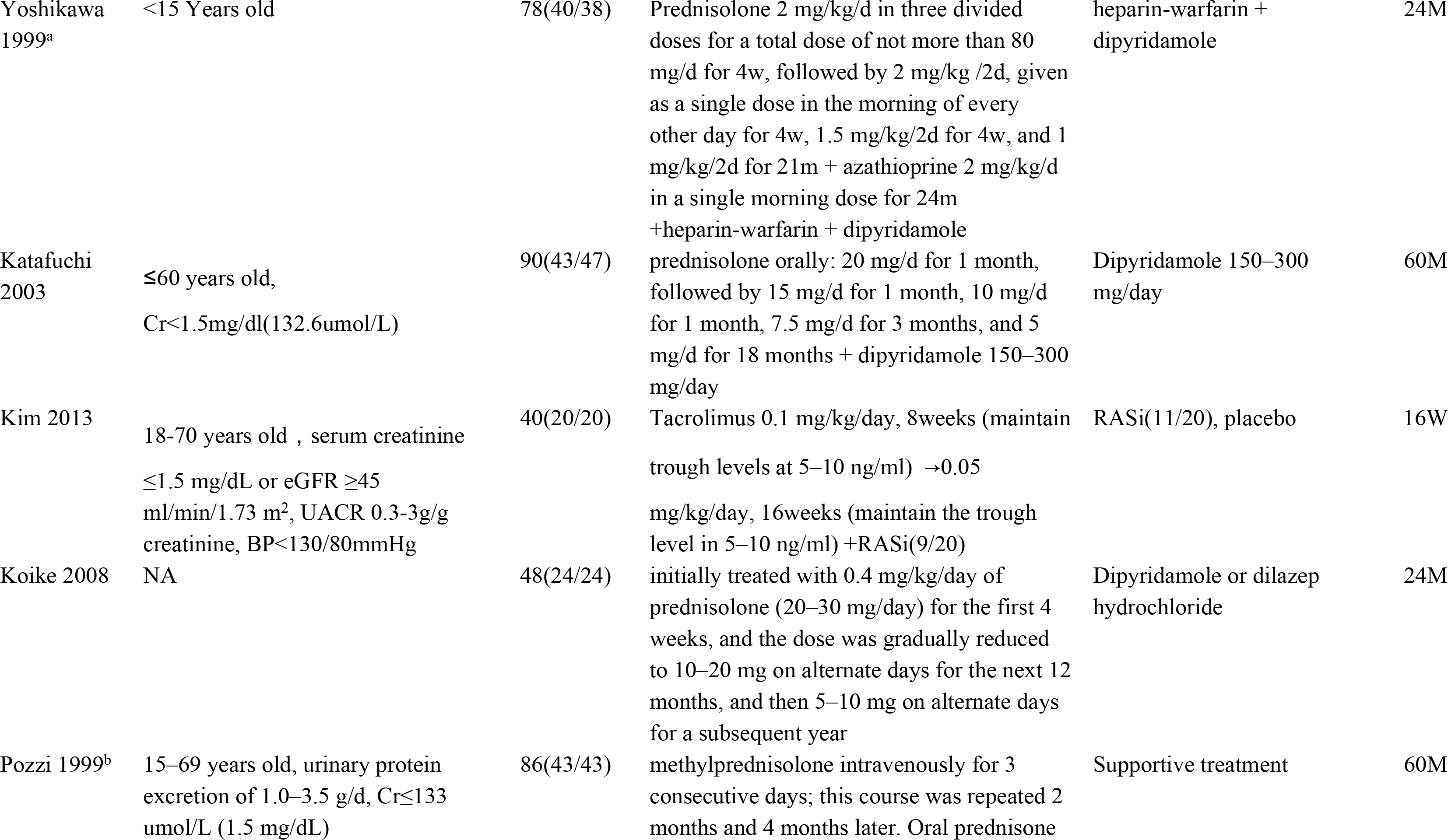
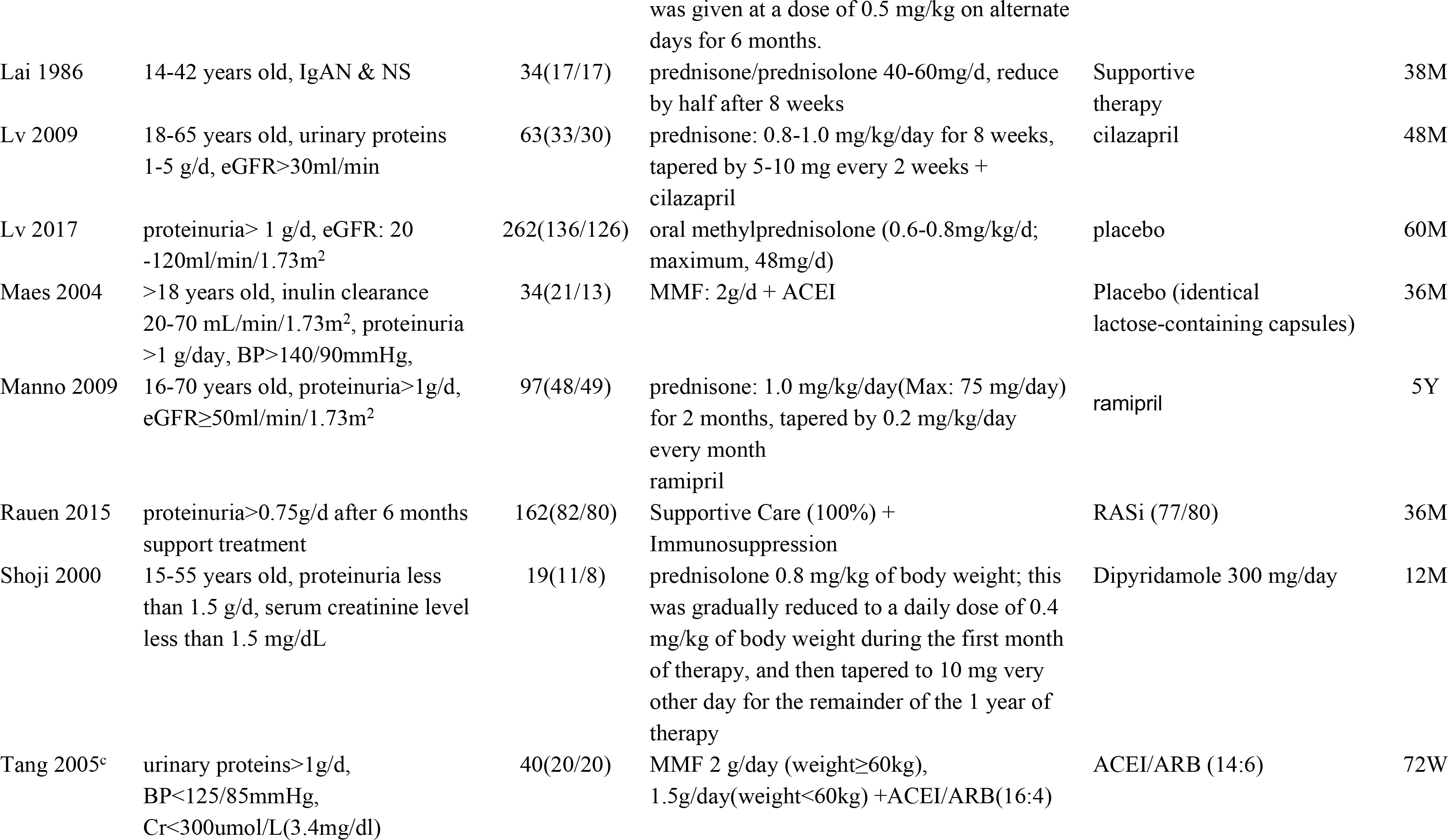
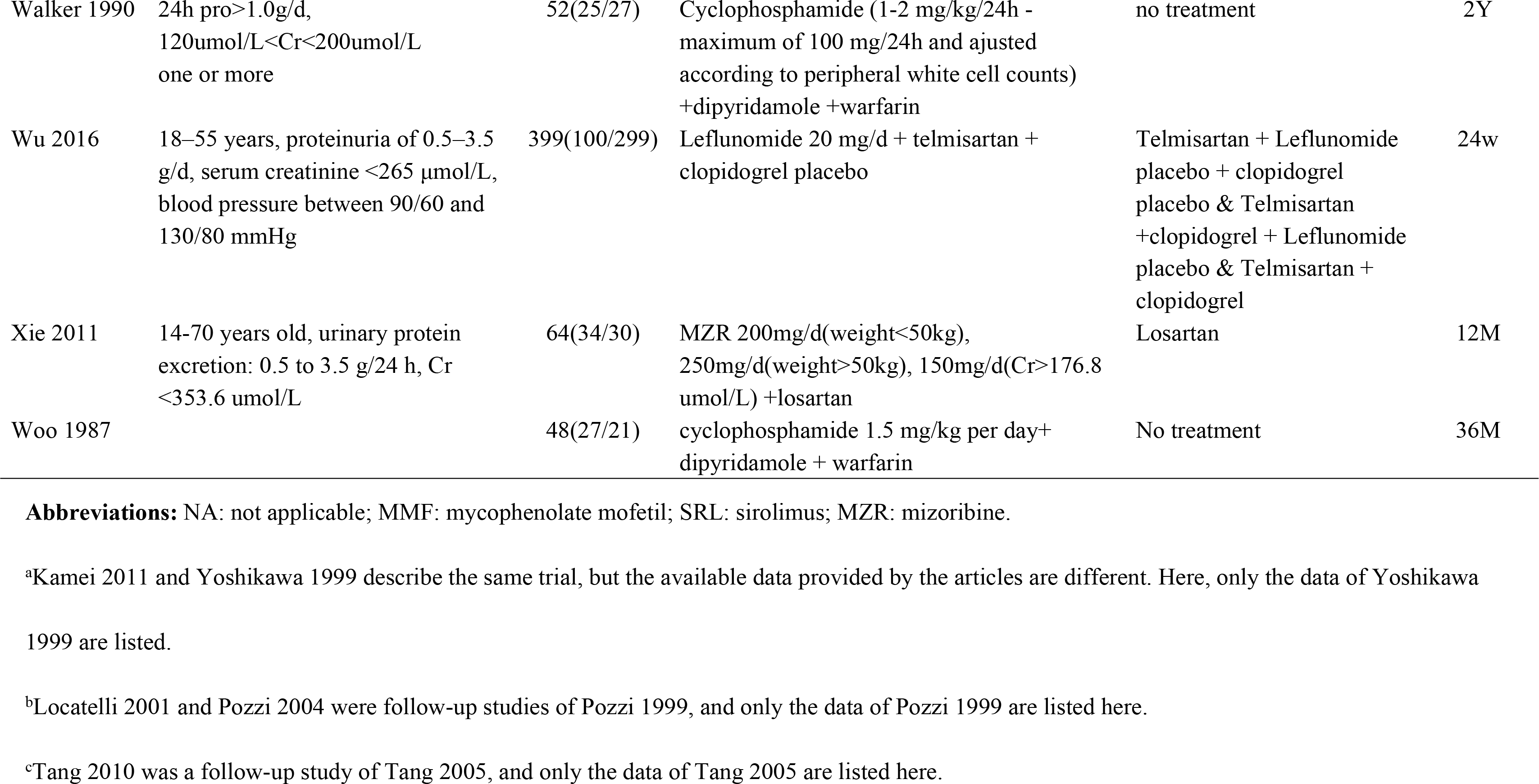
Characteristics of RCTs included in the study

### Quality of trials

By current standards, reporting of key indicators of trial quality was suboptimal. Some studies in particular provided few details on the process of randomization and concealment of allocation. Only six studies were double-blinded trials. Seven studies used an open-label design. The bias and overall risk diagrams of the included studies are presented in Figure 2.

**Fig. 2.**
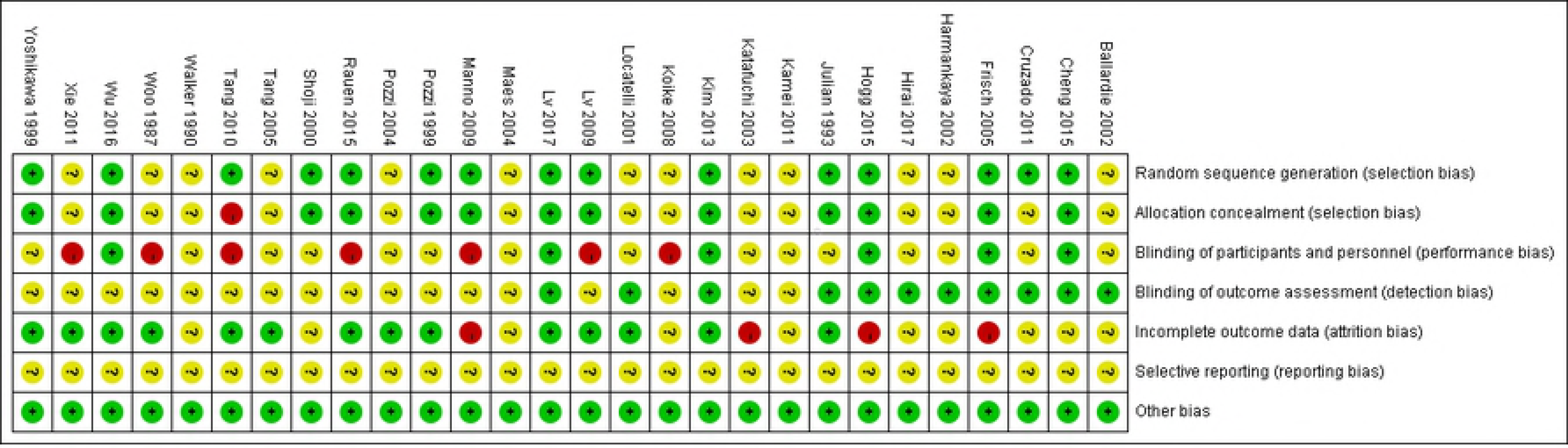
Risk of bias graph.

### Effects on proteinuria

The difference in the means of urinary protein excretion between end of treatment and baseline was significantly lower in the steroid group than in controls (five trials (17, 21-23, 31), 222 patients; WMD −0.51, 95% CI −0.73 to −0.28, with a fixed-effects model; WMD −0.70, 95% CI −1.2 to −0.20, with a random-effects model; I^2^=58%; Fig. 3). After removing Lai (22), heterogeneity I^2^ changed to 0.

**Fig. 3.**
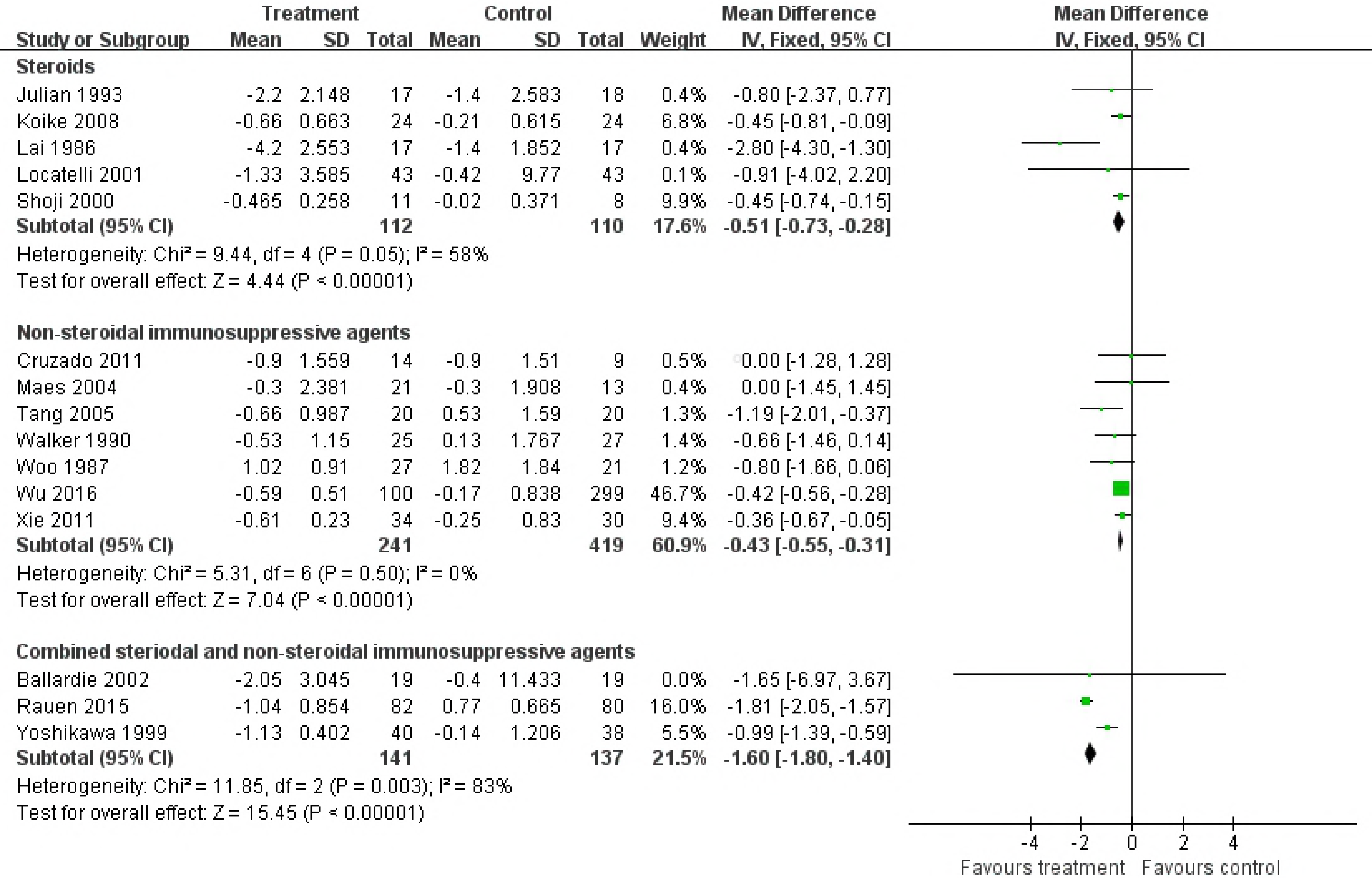
Effects of immunosuppressive agents on proteinuria in patients with IgAN. CI, confidence interval.

Patients receiving NSI alone showed a more significant reduction of urinary protein excretion after treatment compared to controls (seven trials (12, 26, 32, 34-37), 660 patients, WMD −0.43, 95% CI −0.55 to 0.31, with a fixed-effects model; WMD −0. 43, 95% CI −0.55 to −0.31, with a random-effects model; I^2^=0; Fig. 3).

With the S&NSI treatment approach, patients had a more significant reduction of urinary protein excretion after treatment compared to controls (three trials (10, 30, 38), 278 patients, WMD −0.16, 95% CI −1.8 to −1.4, I^2^=83%, with a fixed-effects model; WMD −1.42, 95% CI −2.18 to −0.66, I^2^=89%, with a random-effects model; Fig. 3). After removing Yoshikawa (38), heterogeneity I^2^ changed to 0.

TSAs of steroids, NSI, and S&NSI all indicated that the cumulative z curve crossed both the conventional boundary and the trial sequential monitoring boundary (Fig. 4).

**Fig. 4.**
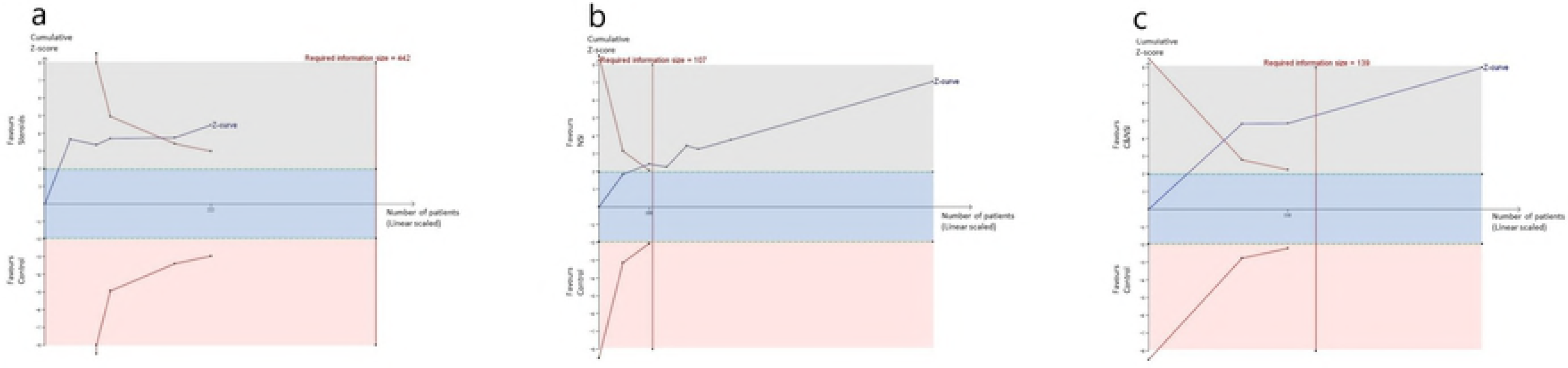
Trial sequential analyses of proteinuria. a) Five comparisons between steroids and controls. b) Seven comparisons between NSI and controls. c) Three comparisons between S&NSI and controls.

### Effects on renal function and renal survival

#### Creatinine

There were no statistically significant differences in creatinine changes between baseline and end of treatment between immunosuppressive treatment and control groups (nine trials (11, 13, 17, 19, 20, 22, 26, 31, 34), 420 patients, WMD −0.03, 95% CI −0.11 to 0.15, with a fixed-effects model; WMD −0.03, 95% CI −0.11 to 0.05, with a random-effects model; I^2^=0%; Fig. 5).

TSAs of nine comparisons illustrated that the cumulative z curve did not cross the conventional boundary or the line of required information size, indicating that the evidence was insufficient. Therefore, further trials are required.

**Fig. 5.**
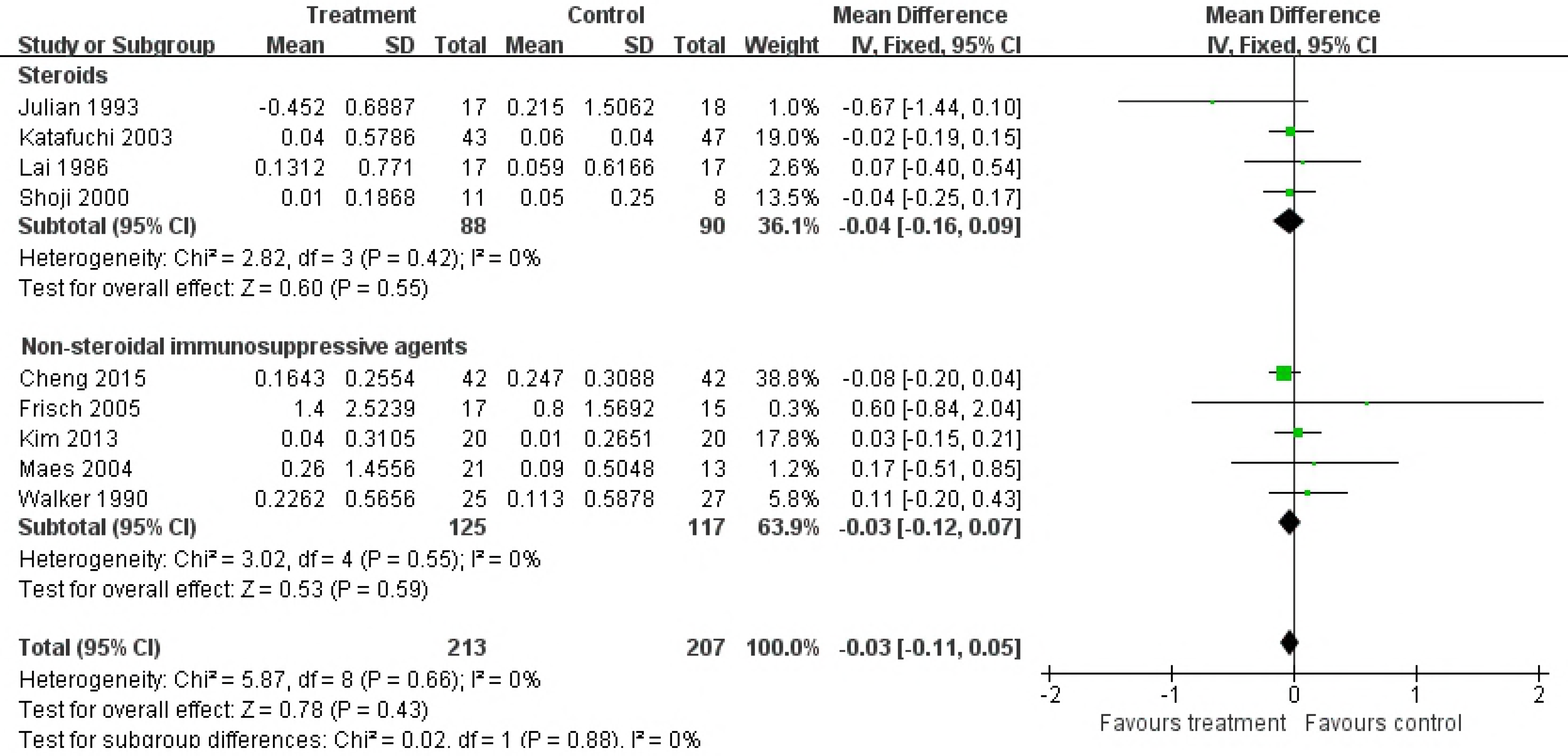
Effects of immunosuppressive agents on creatinine levels in patients with IgAN. CI, confidence interval.

#### eGFR

The differences in the means of eGFR between end of treatment and baseline were significantly higher in the NSI group than in controls (five trials (16, 20, 25, 36, 37), 817 patients; WMD 5.17, 95% CI 3.18 to 7.16, with a fixed-effects model; WMD 5.17, 95% CI 3.18 to 7.16, with a random-effects model; I^2^=0%; Fig. 6). TSAs of five comparisons indicated that the cumulative z curve crossed the conventional boundary, but did not cross the trial sequential monitoring boundary.

However, when the steroid and S&NSI groups were added, there were no significant differences in eGFR changes in immunosuppressive treatment compared to controls (seven trials (16, 20, 25, 30, 31, 36, 37), 998 patients, WMD 0.26, 95% CI −0.03 to 0.56, with a fixed-effects model; WMD 2.52, 95% CI −0.49 to 0.53, with a random-effects model; I^2^=76%; Fig. 6). TSAs of seven comparisons indicated that the cumulative z curve did not cross the conventional boundary or the line of required information size.

**Fig. 6.**
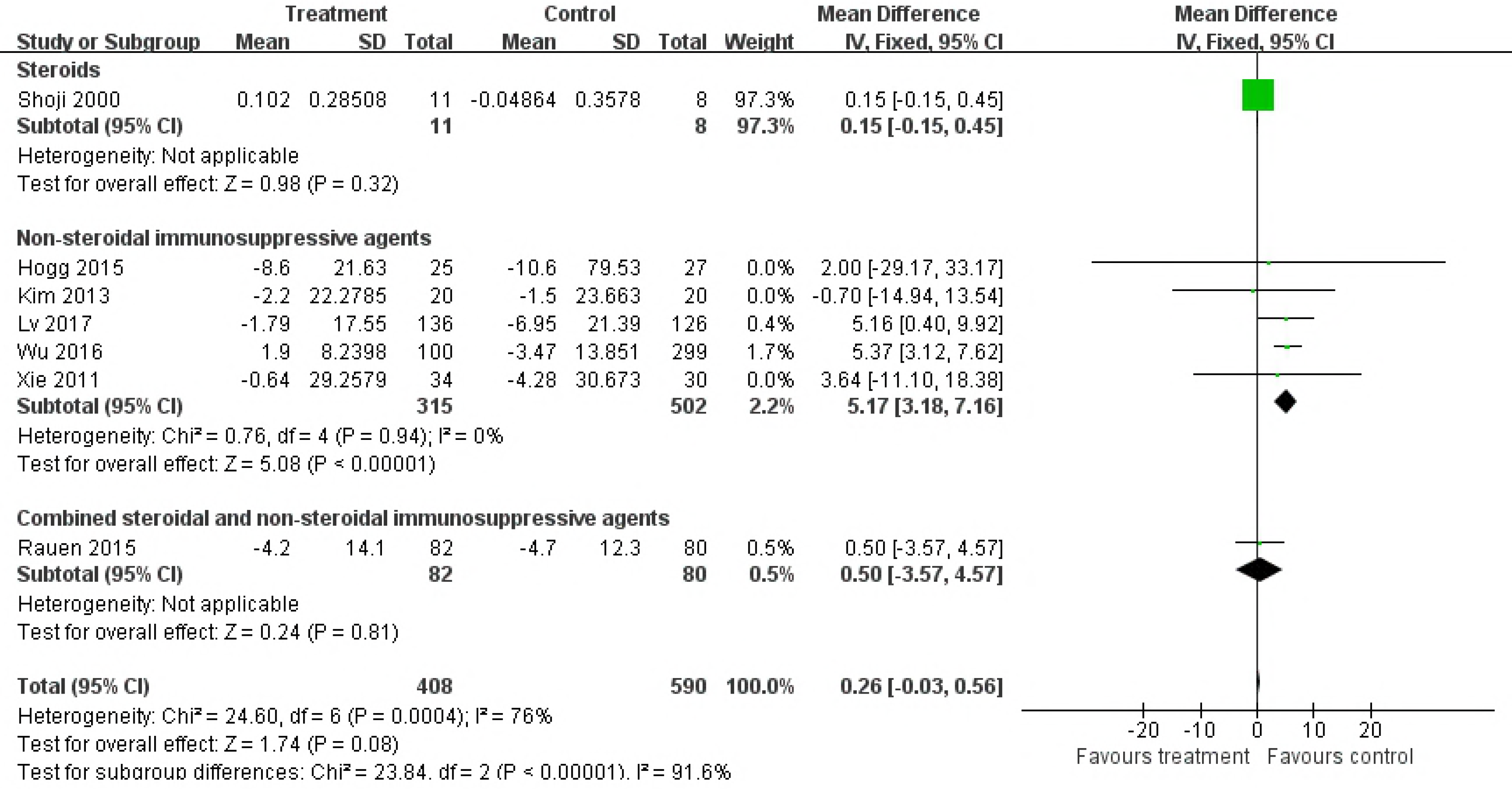
Effects of immunosuppressive agents on estimated glomerular filtration rate in patients with IgAN. CI, confidence interval.

#### ESRD

There was a lower risk of reaching ESRD in the immunosuppressive treatment group than in controls (12 trials (13, 17-19, 24-28, 30, 33, 34), 1031 patients; RR 0.51, 95% CI 0.33 to 0.08, with a fixed-effects model; RR 0.55, 95% CI 0.33–0.90, with a random-effects model; I^2^=8; Fig. 6). These analyses were dominated by the steroid treatment group (Fig. 7).

TSAs of steroids indicated that the cumulative z curve crossed both the conventional boundary and the trial sequential monitoring boundary.

**Fig. 7.**
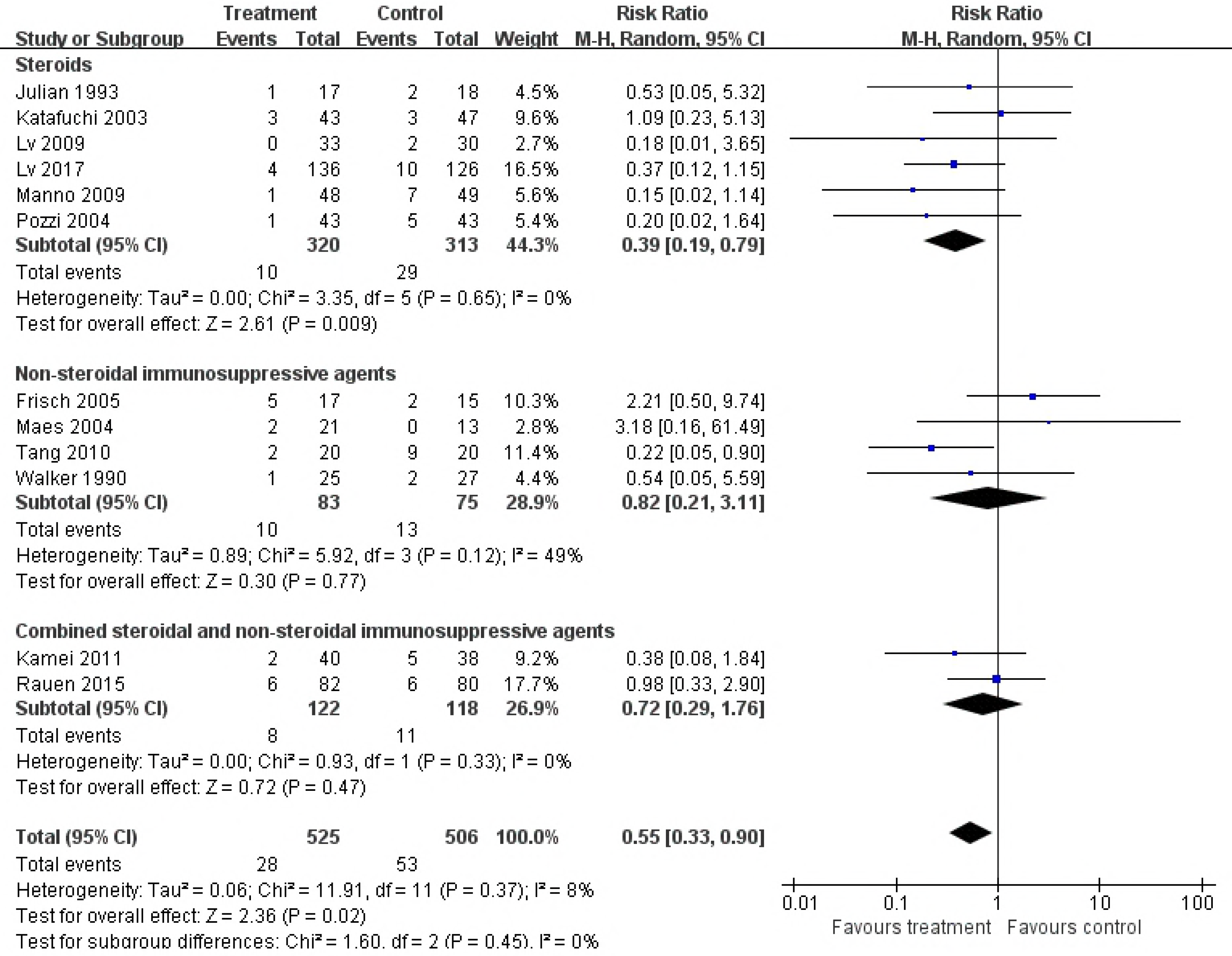
Effects of immunosuppressive agents on end-stage renal disease in patients with IgAN. CI, confidence interval; RR, relative risk.

### Adverse events of treatment

A total of 20 articles reported adverse events during the observation period. The types of adverse events varied widely, and included infection, cardiovascular disease, respiratory disease, hepatotoxicity, and many others; the 12 most commonly reported are listed in Table 2. As the number of infections reported in Rauen (30) was greater than the total number, RR could not be calculated for infections. TSAs of infection, gastrointestinal disease, hematological disease, dermatological disease, impaired glucose tolerance or diabetes mellitus, and hyperkalemia indicated that the cumulative z curve crossed the conventional boundary but did not cross the trial sequential monitoring boundary. In addition, TSAs of the other six diseases indicated that the cumulative z curve did not cross the conventional boundary or the line of required information size.

**Table 2.**
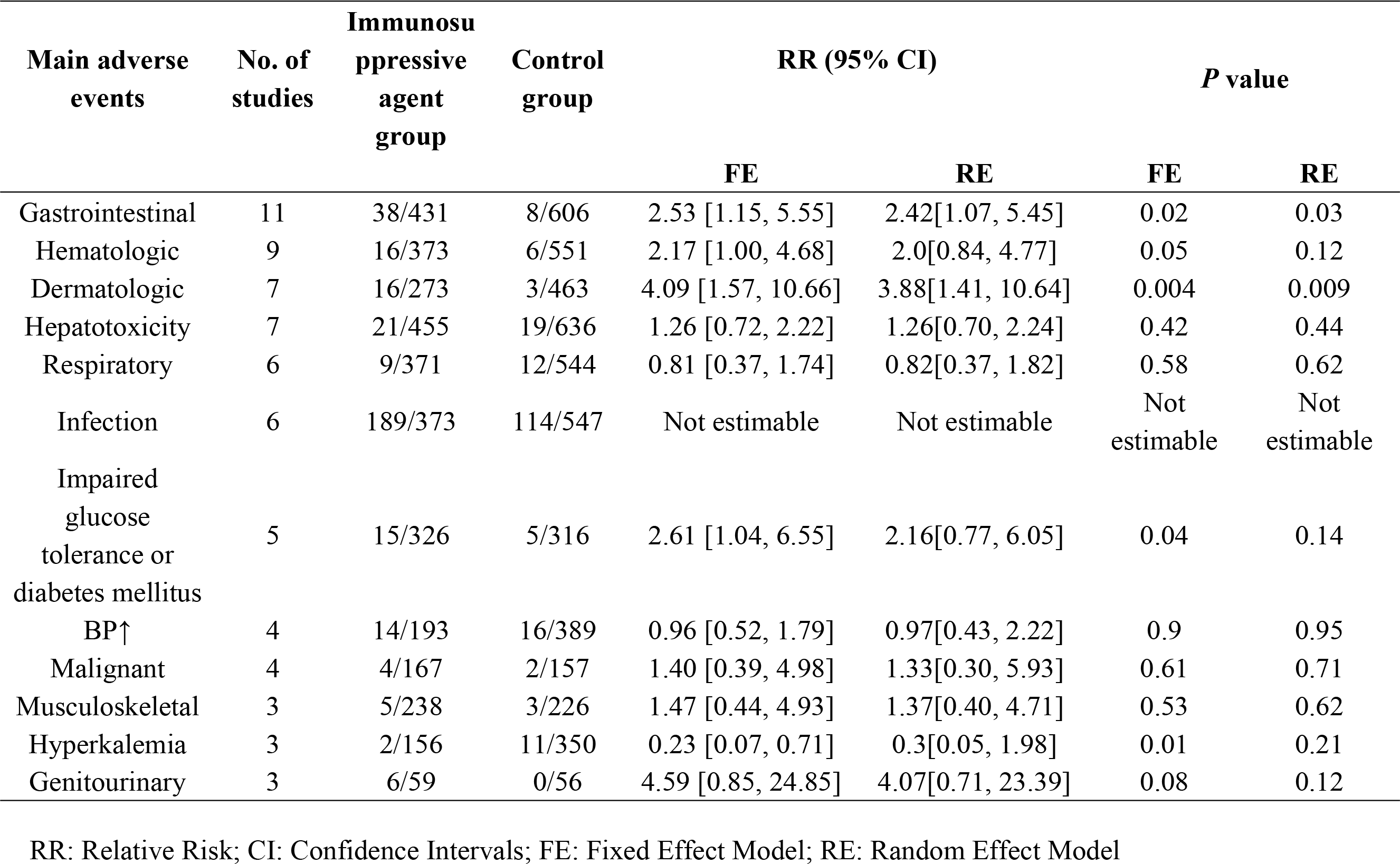
Main adverse events reported in the included RCTs

## Conclusions

Farnsworth (39) and Barnett (40) first used corticotropin between 1949 and 1950 for the treatment of lipoid nephrosis, which is now known as minimal change disease or childhood nephrotic syndrome. Chasis *et al*. (41) used nitrogen mustard to treat chronic glomerulonephritis and achieved good initial results, thus pioneering the use of immunosuppressive agents for the treatment of nephropathy. Immunosuppressive agents have been used for the treatment of kidney diseases for about 70 years. However, the outcomes immunosuppressive therapy for IgAN are controversial. Therefore, we included 29 reports published between 1986 and 2017 in a meta-analysis of the efficacy and safety of immunosuppressive treatment and control treatment in IgAN.

### Alleviation of proteinuria

Previous studies have suggested that treatment with steroids or alkylating agents can significantly reduce proteinuria levels in patients with IgAN (42–44). Our meta-analysis also showed that immunosuppressive agents can significantly reduce the level of proteinuria. The levels of proteinuria in groups treated with steroids, NSI, or S&NSI were significantly reduced compared to controls. The heterogeneity of the steroid group was mainly derived from Lai (22), in which the inclusion criterion included nephrotic syndrome. In addition, the heterogeneity of the S&NSI group was mainly derived from Yoshikawa (38), in which the inclusion criterion included age < 15 years. Sequential analyses showed that immunosuppressive agents were effective for relieving proteinuria, and no additional sample size was required.

### Reducing the risk for ESRD

Our results suggest that non-steroidal immunosuppressive therapy may have a positive effect on eGFR. However, sequential analyses suggested that this is still inconclusive and further studies are required for confirmation. In addition, the treatment group showed a greater reduction in the risk for ESRD than the control group, and this effect was mainly due to the steroid treatment group. Sequential analyses showed that steroids could reduce the risk for ESRD without the need for a larger sample size. A relevant study (43) also suggested that high-dose short-course steroid therapy has a significant protective effect on renal function, while a low-dose long-course of steroids does not. Further studies are required to determine whether NSI or S&NSI can reduce the risk for ESRD.

### More adverse events

The use of immunosuppressive agents is often accompanied by side effects. The immunosuppressive therapy group showed significant increases in gastrointestinal, hematological, dermatological, and genitourinary side effects, as well as impaired glucose tolerance or diabetes in this meta-analysis. As the number of infection events reported in the STOP study was too high, even exceeding the total number of patients, it was not possible to calculate the RR value. However, across all studies, the proportion of infections reported was still higher in the immunosuppressive therapy group than in controls. In addition, the TESTING study had to be discontinued because of the excessive number of serious adverse events, mostly infections. By contrast, hyperkalemia was more common in the control group, which may have been related to the application of ACEI and ARB. However, it should be noted that sequential analyses indicated that the statistical results of the above adverse events should be verified by further experiments.

### Strengths and limitations

Our study had several limitations that should be taken into consideration. The results of bias analyses indicated that nearly half of the studies did not explicitly report the methods used for randomization. In addition, few studies used blinded methodologies. The quality of the reports in the literature is unsatisfactory. In addition, there were some differences in the inclusion criteria between each study, such as age, proteinuria level, and renal function, and these confounding factors led to a high degree of data heterogeneity.

In conclusion, immunosuppressants significantly reduce proteinuria and decrease the risk for ESRD but also increase the risk for serious adverse reactions. Therefore, if it is necessary to use immunosuppressive agents, clinicians should evaluate the patient on an individual basis according to their own conditions before treatment. In the course of using immunosuppressive agents, close observation should be carried out to prevent and control complications. In addition, further well-designed and high-quality RCTs are needed to explore the applicability and optimal methods of immunosuppressant treatment.

The English in this document has been checked by at least two professional editors, both native speakers of English. For a certificate, please see: http://www.textcheck.com/certificate/gC26x1

Author contributions statement
Zheng Zhang designed this study and conducted literature retrieval, data extraction, data analysis and article writing.
Yue Yang participated in literature retrieval, data extraction and data analysis. Shi-min Jiang participated in data extraction.
Wen-ge Li guided the research.

## Notes

**Competing interests statement:** There is no conflict of interest in this article.

